# Ecological plasticity explains the distribution of sympatric and allopatric mouse lemurs (*Microcebus* spp.) in northeastern Madagascar

**DOI:** 10.1101/2025.03.24.645043

**Authors:** Dominik Schüßler, Tobias van Elst, Naina R. Rabemananjara, Tahiriniaina Radriarimanga, Stephan M. Rafamantanantsoa, Roger D. Randimbiharinirina, Emmanuel Rasolondraibe, Jasmin Mantilla-Contreras, Ute Radespiel

## Abstract

Species distributions are shaped by complex biotic and abiotic interactions. We studied major ecological drivers of the distribution of four cryptic mouse lemur species (*Microcebus* spp.) in northeastern Madagascar, across sympatric and allopatric ranges. Using structural habitat characteristics, adaptability to habitat degradation, bioclimatic niches, and morphology, we estimated n-dimensional hypervolumes of niche sizes and overlaps. Annual body mass variability was analyzed as an indicator of heterothermy, a potential indicator for ecological plasticity.

*M. jonahi* and *M. simmonsi* were found almost equally often in forest- and fallow-derived habitats (52% and 58%, respectively), while *M. lehilahytsara* predominantly occupied fallow habitats (71%), likely driven by the coexistence with other species. *M. lehilahytsara* exhibited the largest niche and range size, followed by *M. jonahi* and *M. simmonsi*. In contrast, *M. macarthurii* was restricted to a narrow niche and range. *M. jonahi* and *M. simmonsi* displayed significant fat deposition before the austral winter, indicating heterothermy as an adaptation to unpredictable environmental conditions. These species were only found in allopatry, likely due to competition for critical resources like sleeping sites. The smaller *M. lehilahytsara* coexisted with *M. jonahi* and *M. macarthurii* potentially by adjusting its niche to avoid competition with these larger-bodied species.

We hypothesize that heterothermy allowed *M. jonahi*, *M. simmonsi*, and *M. macarthurii* to persist in lowlands during Pleistocene climatic challenges but that it did not promote coexistence. In contrast, *M. lehilahytsara* may have lost lowland distributions due to a lower ability to store fat. Our findings require further testing but provide a plausible explanation for a complex distributional pattern of sympatric and allopatric occurrences.

## INTRODUCTION

The distributions of species are modulated by a multitude of factors, including biotic and abiotic interactions, innate differences in environmental plasticity, evolutionary history and dispersal abilities (Acevedo et al., 2016; Boulangeat et al., 2012; Gaston, 2003; Sexton et al., 2009). Species are found where favorable abiotic conditions, complementary co-occurring species, and ecological accessibility converge (Sexton et al., 2017; Soberón and Peterson, 2005), while their evolutionary history fundamentally determines the areas in reach for dispersal and expansion after speciation (Avise, 2000; Zeisset and Beebee, 2008).

Apart from biogeographic drivers, the ecological niche of a species constrains its distribution (Grinnell, 1917; Sexton et al., 2017) and can be described by an “n-dimensional hypervolume”, spanning a variety of axes of environmental variables (Hutchinson, 1957). Taxa with broader niches tend to occupy wider geographical ranges and coexist with more species due to their ability to persist under diverse conditions and to exploit a wider range of resources (Brown, 1984; Hausharter et al., 2023; Kambach et al., 2019; Slatyer et al., 2013). A special case are cryptic species, which are closely related taxa without much phenotypic distinctiveness (Bickford et al., 2007). They are expected to be restricted by their phenotypic and ecological similarity limiting their potential for coexistence (Cothran et al., 2013; Violle et al., 2011; Vodā et al., 2015).

We aim to investigate the ecological basis of coexistence of four closely related and cryptic mouse lemur species (*Microcebus* spp.) in northeastern Madagascar. The few sympatric occurrences in this genus appear to be driven by two widely ranging species partly coexisting with locally restricted endemics (van Elst et al., 2024; Radespiel, 2016). Coexistence with *M. murinus*, for example, could already be related to a differential use of sleeping sites, food resources, microhabitats and heterothermy (Dammhahn and Kappeler, 2008a; Dausmann and Warnecke, 2016; Radespiel et al., 2003, 2006; Rakotondranary et al., 2011; Rakotondranary and Ganzhorn, 2011; Sehen et al., 2010). It likely colonized the ranges of all species potentially in reach rather recently (Blair et al., 2014, van Elst et al., 2024, Schneider et al., 2010). Mechanisms of coexistence with *M. lehilahytsara*, the other widely ranging species, remain largely unknown. Although the ranges of multiple *Microcebus* species are in reach (e.g., *M. gerpi, M. simmonsi, M. jonahi* and *M. macarthurii)*, *M. lehilahytsara* has only been found in sympatry with the latter two (van Elst et al., 2024, 2025). Furthermore, sympatry with *M. lehilahytsara* is only restricted to specific parts of the distribution of *M. jonahi* and *M. macarthurii* (van Elst et al., 2025; Figure 1). In one particular inter-river system, *M. jonahi, M. simmonsi* and *M. lehilahytsara* can even be found together, but with the latter only occupying higher elevations together with *M. jonahi*, while this species occurs in allopatry at mid-elevations and *M. simmonsi* in allopatry at lower elevation (van Elst et al., 2025; Figure 1). There are no geographic barriers between these occurrences and an elevational gradient is unlikely due to lowland and highland occurrences of all three species (van Elst et al., 2024, 2025). Whether this seemingly unusual distribution pattern of *M. lehilahytsara*, with partial lowland occurrences followed by absences, partial sympatry and partial allopatric occurrences is due to competitive exclusion or reflects other ecological constraints, is unknown so far.

**Figure 1:**
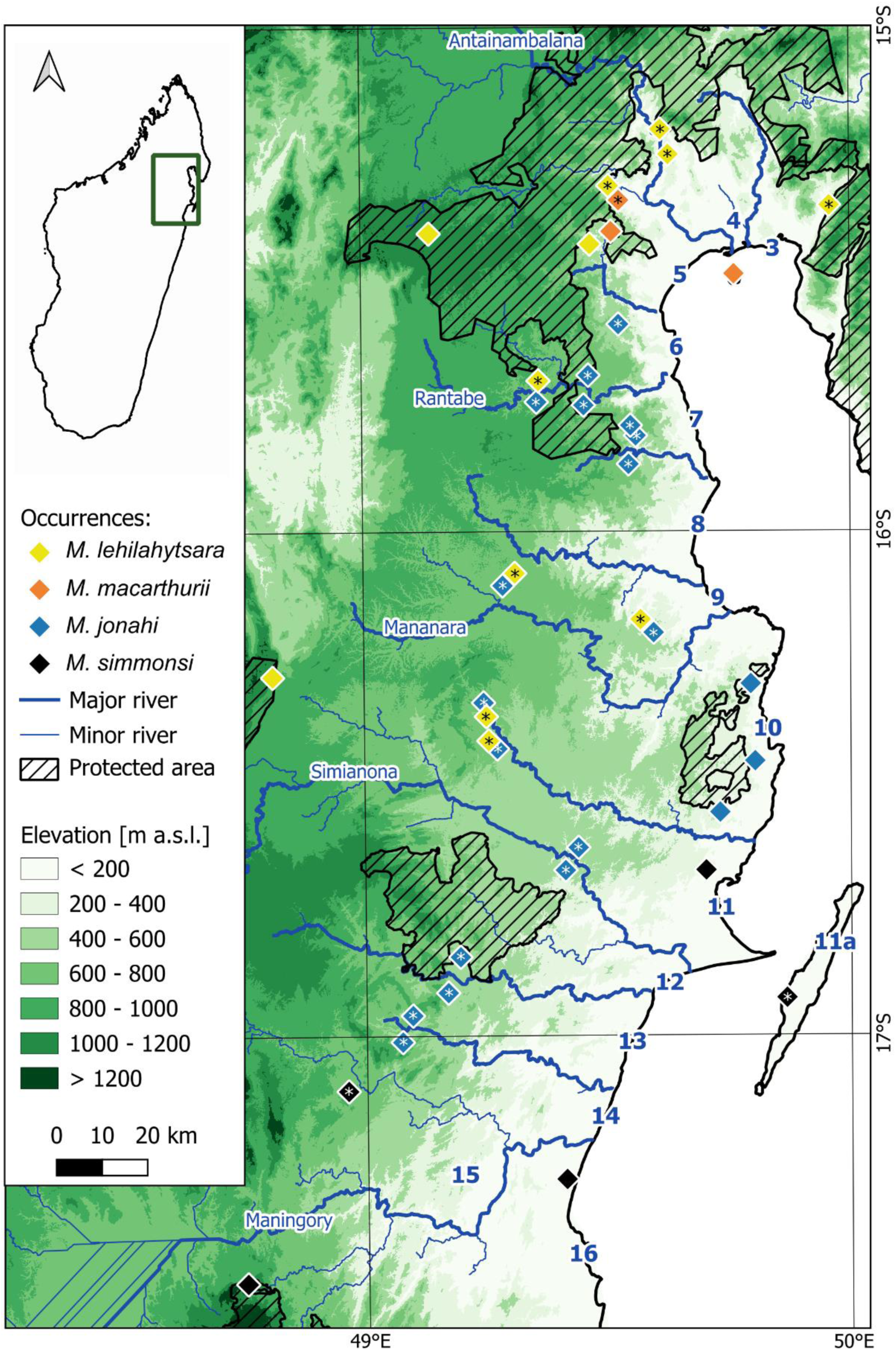
Study sites with species occurrences, supplemented from van Elst et al. (2024). Newly surveyed sites are marked with an asterisk (van Elst et al., 2025). Blue numbers indicate the inter-river systems of the region. Relevant rivers and associated names are given in blue. Elevation (JAXA, 2015), rivers (Schüßler et al., 2025) and protected areas (UNEP-WCMS, 2023) are also illustrated.

We aim to characterize the mechanisms that enable coexistence of these four closely related cryptic species and the interplay between ecological factors that shaped the allopatry-sympatry pattern found today. The following hypotheses shall be tested:

1. Population density of mouse lemurs in general is equal in areas of allopatric and sympatric occurrence due to competition for the same critical resources (principle of density compensation; MacArthur and Wilson, 1967).
2. The partitioning of the multidimensional niche spaces (vegetation structure, landscape-scale degradation, bioclimatic niche, morphology) explain why some species occur in sympatry and others in allopatry.
3. Heterothermy, as indicated by an annual body mass variability, allows for temporal niche shifts during times of resource scarcity (Blanco et al., 2018; Dausmann and Warnecke, 2016).

## METHODS

### Study region

Northeastern Madagascar is a tropical rainforest region (Beck et al., 2018) characterized by primary rainforests, degradation stages thereof, and deforested areas in proximity to major roads and larger cities (Schüßler et al., 2020b). Shifting cultivation for staple crop and agroforestry for cash crop production are the major land use types (Llopis et al., 2019; Martin et al., 2023; Zähringer et al., 2015). The region is ecologically structured by large rivers that represent distributional barriers for some but not all species (Schüßler et al., 2025). We therefore refer to areas with between these major rivers as inter-river systems (IRS, Figure 1). The region harbors four different mouse lemur species, *Microcebus jonahi*, *M. lehilahytsara*, *M. macarthurii*, *M. simmonsi*, that occur both in allopatry and in sympatry (Figure 1; van Elst et al., 2025). Data collection was conducted at 23 sites across the entire region between IRS 3 and 15 (Figure 1) during August - September 2017, August - November 2019, September - December 2021 and March - May 2022. Sampling sites were all situated outside of the protected area network within village-owned forests and fallows and visited for 10-14 days each.

### Mouse lemur surveys

We installed 109 standardized line transects (mean length: 1.25 km, range: 0.20-4.20 km depending on availability of pathways and similarity of habitat; total effort: 310.44 km) that were surveyed 1-5 times by a team of 3-6 observers (walking speed 1 km/h). Our transects crossed six vegetation types with the highest survey effort in selectively and highly logged primary rainforest (Table 1; Table S1). Nocturnal surveys were conducted during the first activity peak of mouse lemurs (18h00–22h00). Sightings were noted by taking position data (GPS coordinates, Garmin GPSMAP 64s), date and time, perpendicular distance estimated using a laser distance meter (Bosch Professional GLM 40), and the number of individuals. Mouse lemurs were detected with head lamps by eye reflections and confirmed using a powerful hand torch. Species assignment was not possible during surveys due to their cryptic morphology, but was clarified during parallel capture and genomic identification (van Elst et al., 2025; see below). Additional ad libitum GPS coordinates were taken at any time a mouse lemur was sighted.

**Table 1:**
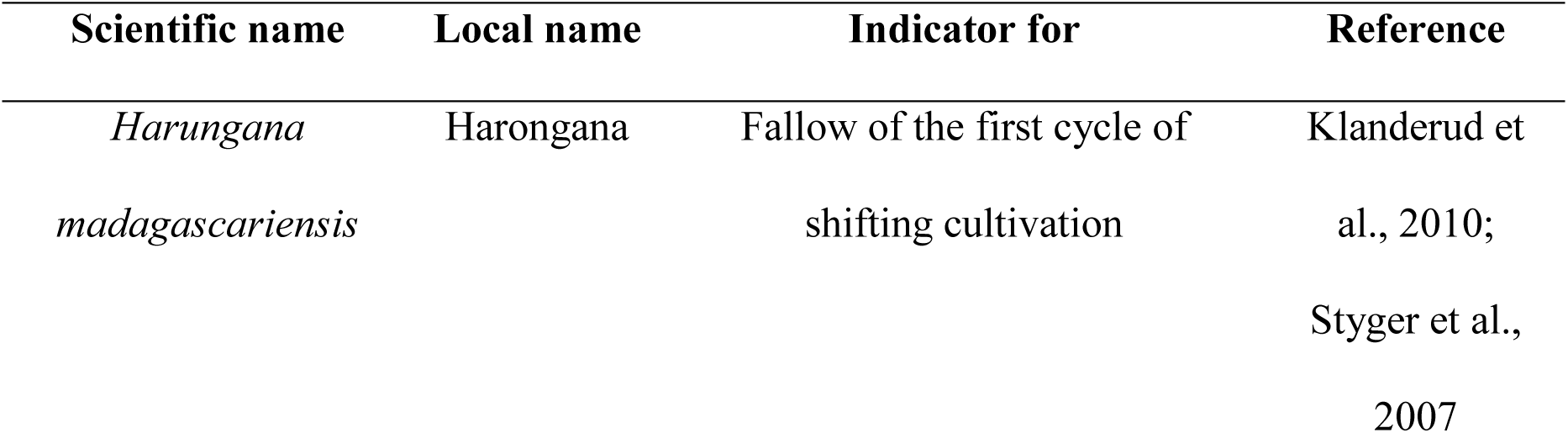
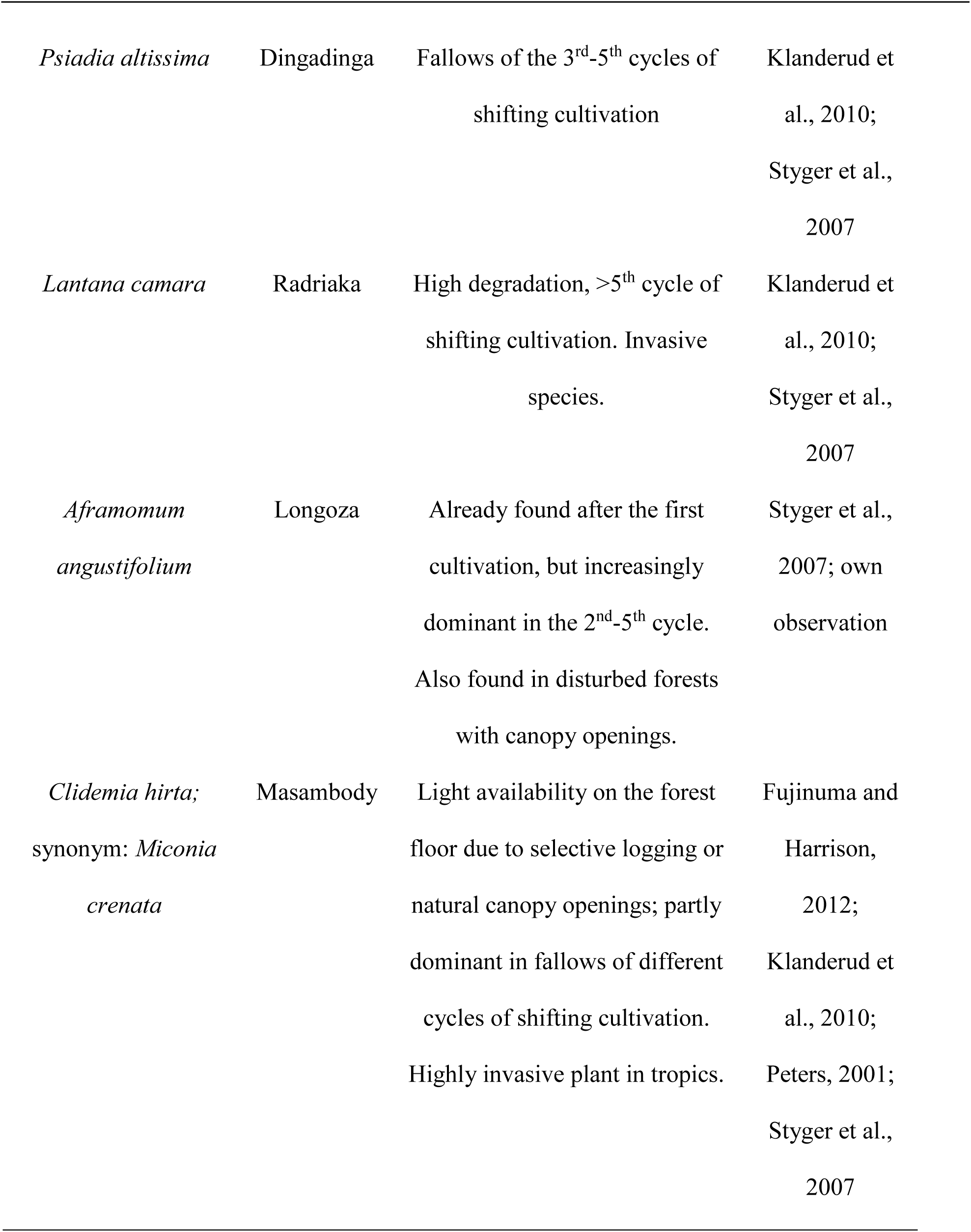
Indicator species for land use history and habitat structure in northeastern Madagascar that are relevant within mouse lemur habitats.

### Abundance estimation

We calculated overall mean mouse lemur encounter rates per transect km (Buckland et al., 2001) and compared these across the different habitat types (analysis of variance, ANOVA, with Tukey HSD post-hoc tests) and areas with sympatric and allopatric occurrences (t-tests). Mouse lemur population densities were estimated as individuals per km² (ind./km²) following distance sampling protocols using the “distance” package (v.1.0.9; Buckland et al., 2001; Miller et al., 2017) in R and RStudio (Posit Team, 2024; R Core Team, 2024). Data were truncated at 35.0 m and the best detection function was determined by the lowest AIC value (Akaike information criterion; Burnham and Anderson, 2002; Miller et al., 2019). The fit of the detection function was checked via visual interpretation of quantile-quantile plots and χ^2^-goodness-of-fit tests (Buckland et al., 2001; Miller et al., 2019). Density estimates were separately calculated for sympatric and allopatric areas. Further estimates were given for *M. jonahi* in allopatry, which was not possible for the other species due to limited data.

### Habitat structure at sighting locations

Structural habitat data were collected for about 90% of all sighting locations. We surveyed the vertical vegetation structure by estimating coverage in five height strata (<0.5, 0.51−2.0, 2.1−5.0, 5.1−10.0, >10.1 m) within a 5.6 m radius of each sighting location (equivalent to a 100 m² plot), using the LONDO scale (Londo, 1976). This scale facilitates the estimation of vegetation coverage using bins (e.g., >1.0% coverage, 1.1−3.0%, 3.1−5.0%, 5.1−10.0%, 10.1−15.0%, etc.). Coverage for each stratum was estimated by an experienced observer, with frequent validation by other team members (Schüßler et al., 2023b). For analysis, the intermediate coverage value within each bin was used (e.g., 10-15% coverage had a bin value of 13). Overall coverage of eight indicator plant species of land use history was further estimated (Table 1).

In addition, we categorized each plot by its degradation stage (Table 2) and further subdivided these based on their land use history into forest-derived and fallow-derived habitats (Martin et al., 2020b; Styger et al., 2007). As we are in favor of abandoning the primary-secondary forest dichotomy (Pain et al., 2020), we define our habitat categories as primary (or old growth) forests of different degradation stages compared to vegetation re-growing on land that was at least once burned for shifting cultivation purposes (Table 2).

**Table 2:**
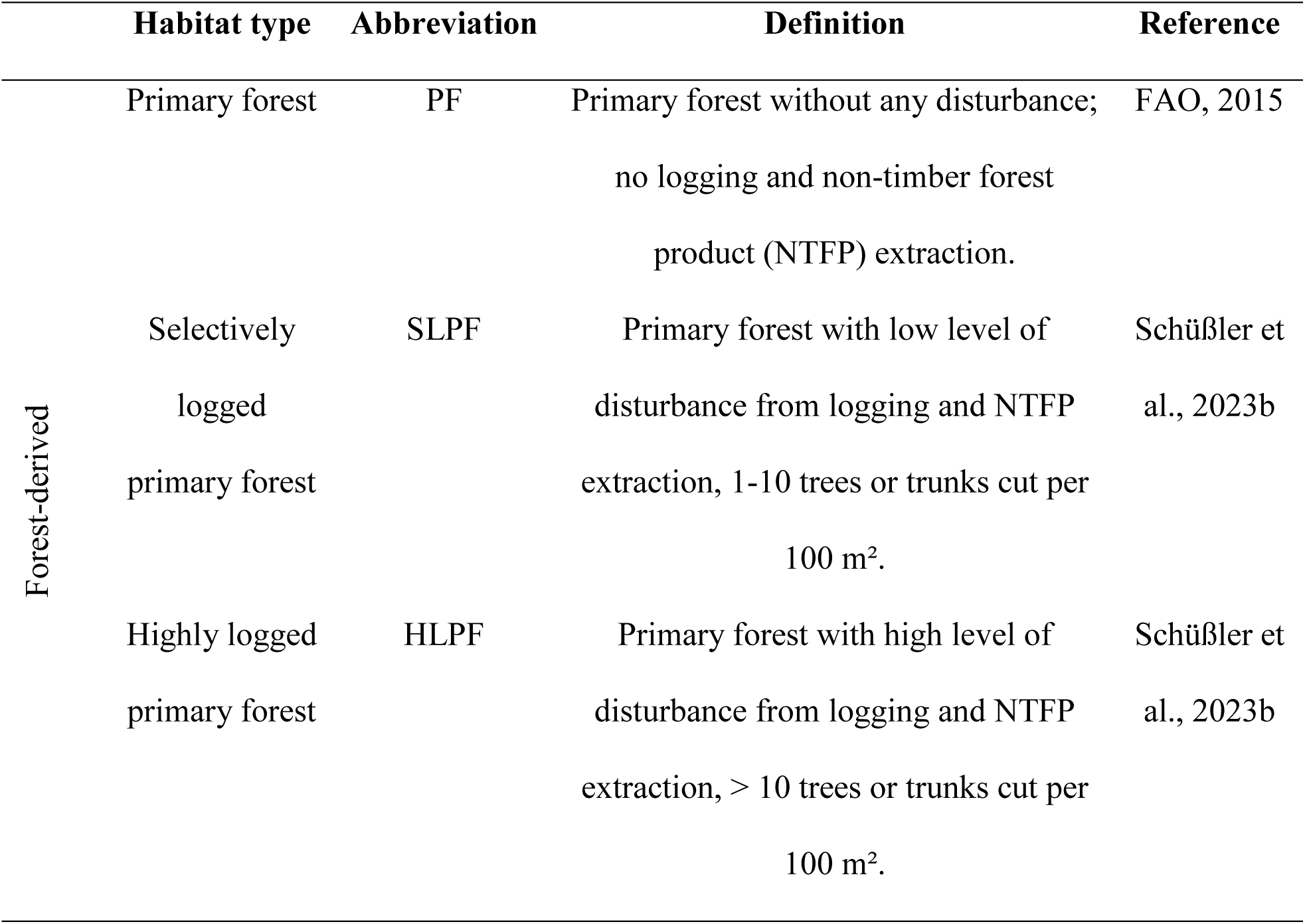
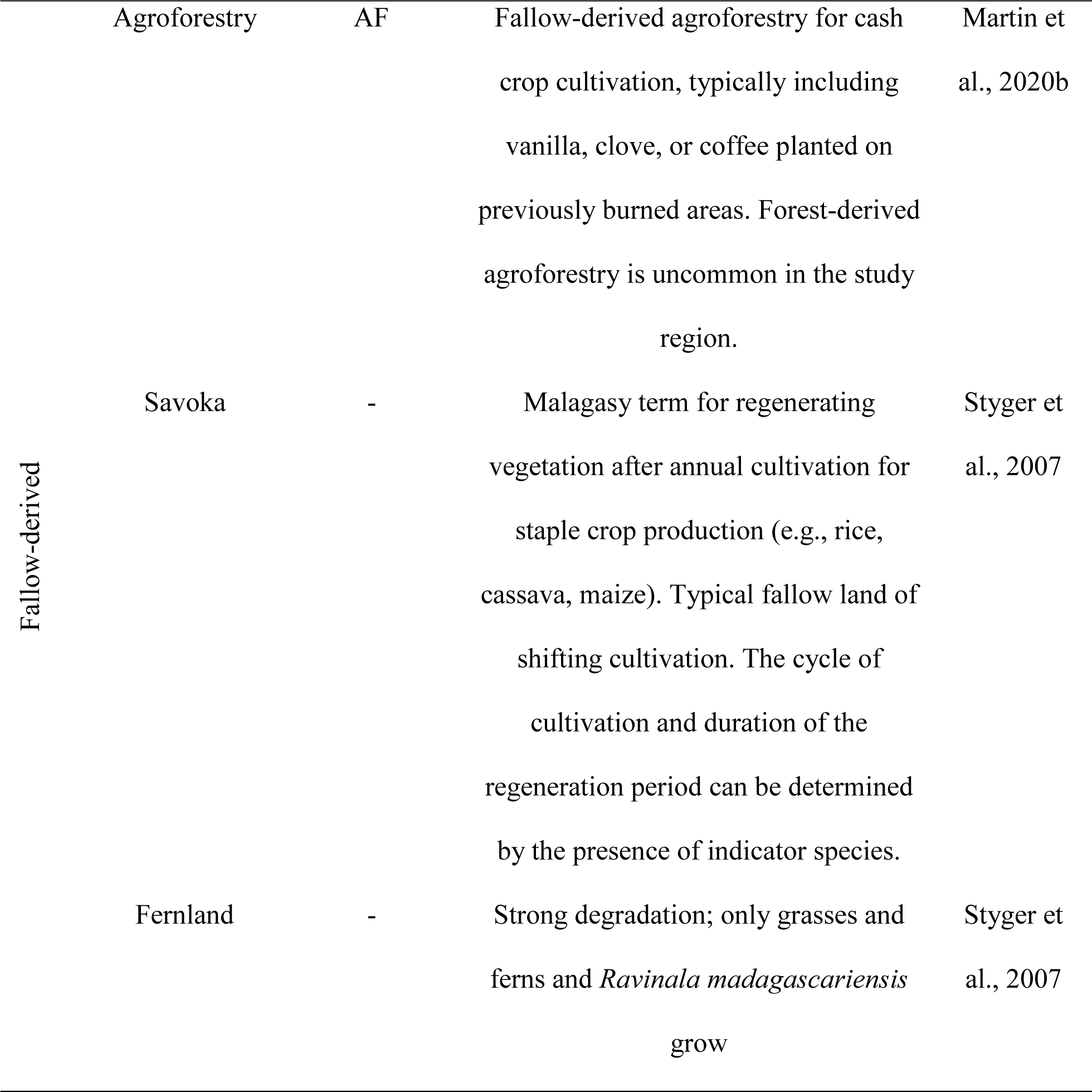
Habitat type definitions based on the recent land use practices.

Species level occurrence proportions in forest- and fallow-derived habitats and between allopatric and sympatric populations were further tested (Poisson test, Fisher’s exact tests with Bonferroni correction after multiple testing). Vegetation data (i.e., coverage in strata and of indicator species) were checked for high correlations r ≥│0.70│ (“psych” package v.2.4.6; Revelle, 2024), which did not exist. Data processing and visualization were conducted using R packages “readxl” (v.1.4.3 Wickham and Bryan, 2023), “dplyr” (v.1.1.4; Wickham et al., 2023), and “ggplot2” (v.3.5.1; Wickham, 2016).

### Morphological measurements

In a parallel study, van Elst et al. (2025) captured mouse lemur individuals from the study populations to genomically identify species and to determine allopatric and sympatric distributions. Morphological data of each captured mouse lemur was also taken and included 13 different variables characterizing ear, head, hind limb and body proportions. Based on this data (supplemented from the literature; Schüßler et al., 2023a), we documented the annual body mass variability per sex and species and characterized the morphological trait axis in the niche differentiation analysis (see below). All procedures adhered to ethical standards of field primatology (ASP, 2001; Riley et al., 2014).

### Multidimensional niche differentiation

We compiled data for four trait axes characterizing (1) vegetation structure, (2) landscape-scale anthropogenic pressure on forest resources, (3) bioclimatic conditions, and (4) morphological traits (Table 3). The data were spatially filtered for bioclimatic variables to exclude duplicate records (Boria et al., 2014). Due to intermediate sample sizes and high dimensionality of the data, we transformed data for each of the four trait axes using a principal component analysis with prior z-score standardization. We used the first three principal components (Blonder et al., 2017; Junker et al., 2016; Lu et al., 2021), which explained at least ≥ 69.45% of the variance in the data (Table S2-S6).

**Table 3:** The four trait axes of niche characterization with definition and variables used.

|  | Definition | Variables | Note |
| --- | --- | --- | --- |
| Vegetation structure | Structural microhabitat characteristics derived from sighting and capture locations, based on the vegetation plots | Coverage in all strata and of indicator species as described above | Non-genotyped individuals in sympatric areas were excluded |
| Landscape-scale degradation | A proxy for adaptability towards modified habitats | Forest density (in 2018) and deforestation density (1990-2018) within a radius of 5 km around the sighting, human population density per km<sup>2</sup>, and aerial distance from sighting to the nearest village | Sightings were supplemented from Miller et al. (2018); variable data from Rakotoarisoa et al. (2024) and Schüßler et al. (2020b) |
| Bioclimatic niche | The bioclimatic niche space was derived from eight bioclimatic variables relevant for mouse lemur ecology (van Elst et al., 2024; Kamilar et al., 2016) | Isothermality, temperature seasonality, maximum temperature of warmest month, minimum temperature of coldest month, annual precipitation, precipitation seasonality, precipitation of wettest and driest quarter | Bioclimatic data from the CHELSA v2.1 database (Karger et al., 2017, 2018); occurrence were supplemented from van Elst et al. (2024). For *M. lehilahytsara*, only records from the study region were used, but not its entire range. |
| Morphology | The morphological variability of the mouse lemur species | Ear length, head length and width, interorbital distance, body and tail length | Data was quality filtered to obtain the variables with the lowest number of missing data and highest inter-observer reliability across all species (Schüßler et al., 2023a) |

We calculated the overlap of n-dimensional hypervolumes across the four trait axes using the “hypervolume” package v.3.1.4 (Chen et al., 2024; Blonder, 2014, 2017). As the hypervolume calculation involves probabilistic components which depend on sample sizes, we first determined the minimum sample size per trait axis and used it for random subsampling. To account for data variability, we repeated the subsampling 100 times and calculated mean values for the size of the hypervolumes and overlaps (Blonder et al., 2017; Chen et al., 2024).

## RESULTS

### Population density and abundances

We recorded a total of 460 mouse lemurs during surveys, 220 in allopatry and 240 in sympatry. Population density was significantly higher in sympatry (t-test: t = - 5.471, df = 367.60, P < 0.001), with 175.0 ind./km² (standard error = 16.3, 95% confidence interval: 145.1-211.2 ind./km²; Table S7, incl. density estimates for *M. jonahi* in allopatry) compared to 72.5 ind./km² (standard error = 8.8, 95% confidence interval: 56.9-92.3 ind./km²) in allopatry.

Mouse lemurs were not equally abundant across habitat types. Mean encounter rates differed significantly (ANOVA: F(4, 99) = 4.589, P = 0.002) with PF having the lowest, HLPF and AF the highest rates (Figure 2, Table S8). While mean encounter rates were higher in sympatry than in allopatry in all vegetation types (Figure 2), this contrast was significant only in SLPF (t-test: t = - 3.608, df = 9.699, P = 0.005; not tested for AF due to small sample size; Figure 2; Table S9). Notably, occurrences in fallow-derived habitats were only regularly noted within a distance of up to 600 m from the forest edge, but not further away from it (SI, Figure S1).

**Figure 2:**
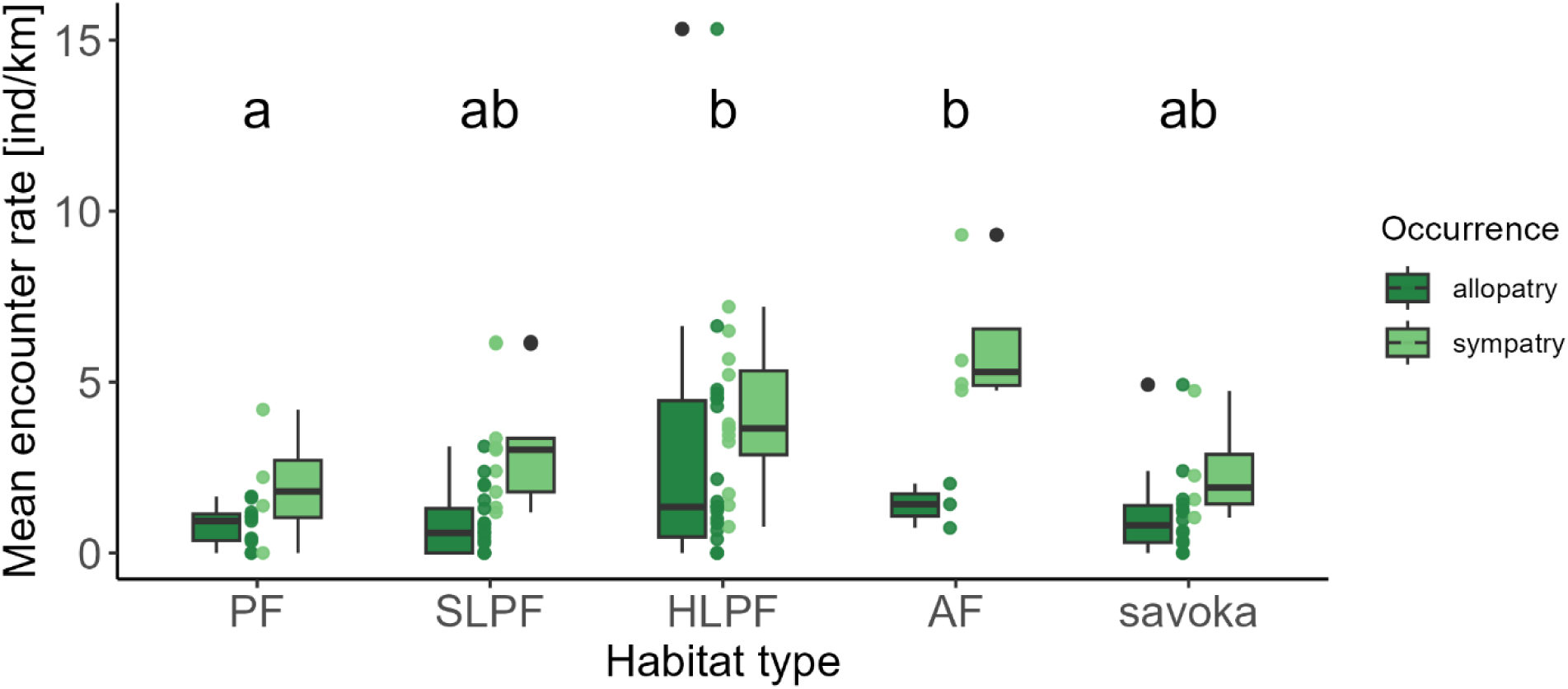
Mean encounter rates of mouse lemurs in different habitat types in sympatry (light green) and allopatry (dark green). Letters above the boxes are based on an analysis of variance with a Tukey HSD post-hoc test of habitat types (without subdivision into allopatric/sympatric occurrences; Table S8). Abbreviations: PF = primary forest, SLPF = selectively logged PF, HLPF = heavily logged PF, AF = agroforestry, savoka = fallow land. Boxplots represent medians with interquartile range (IQR: 25th-75th percentile), whiskers (minimum/maximum values) and outliers (values >1.5*IQR further away from the IQR itself). Individual data points are shown as jitter for the respective dataset next to the respective box plot.

### Niche partitioning in allopatry and sympatry

*Microcebus jonahi* and *M. simmonsi* were almost equally often found in forest-derived (PF, SLPF, HLPF) and fallow-derived habitats (AF, savoka; Figure 3; P ≥ 0.242, Table S10). In contrast, *M. lehilahytsara* was significantly more often observed in savoka (Poisson test, Fisher exact test: P = 0.001; Table S10, S11). *M. macarthurii* only occurred in one inter-river system and in sympatry with *M. lehilahytsara*. Only five individuals were captured from locations that likely promoted their capture, i.e., in agroforestry and savoka, biasing the data (Figure 3). When occurring in sympatry, *M. lehilahytsara* was more likely to occur in fallow-derived habitats than in allopatry (proportions of 0.84 and 0.50, respectively; Fisher exact test: P = 0.009), which was not the case for *M. jonahi* (P = 0.512; Table S12).

**Figure 3:**
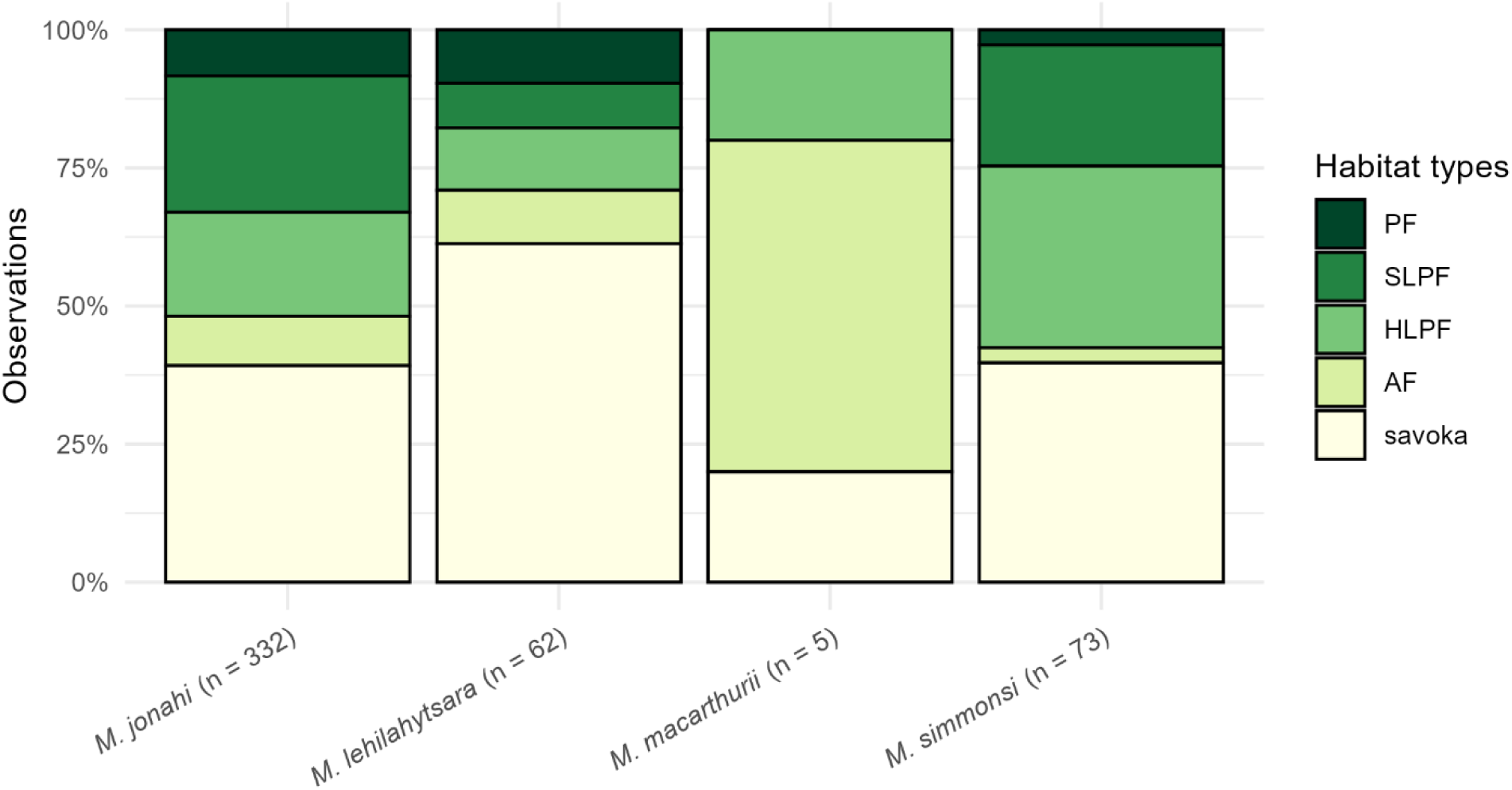
Sighting proportions of *Microcebus* spp. in different habitat types. Abbreviations: PF = primary forest, SLPF = selectively logged PF, HLPF = heavily logged PF, AF = agroforestry, savoka = fallow land. The low sample size for *M. macarthurii* must be treated with caution. PF, SLPF, HLAF are forest-derived habitat types, AF and savoka are fallow-derived.

Niche space size (i.e., hypervolume sizes) differed between *M. jonahi* and *M. lehilahytsara*, with the latter having larger niche spaces along all trait axes (Figure 4). However, there was a variable degree of overlap, particularly in vegetation structure. Compared to *M. simmonsi*, *M. lehilahytsara* showed the same relationships, with *M. simmonsi* having much smaller overall niche spaces than all other species. Except for vegetation structure, overlaps were only marginal (Figure 4). The morphology axis could not be included for *M. simmonsi* due to low sample size. *M. jonahi* and *M. simmonsi* had the highest overlap in vegetation structure, but only marginal overlaps along the other trait axes. *M. jonahi* exhibited a higher degree of niche variability than *M. simmonsi* in general. Similar analyses were not possible for *M. macarthurii* due to small sample size.

**Figure 4:**
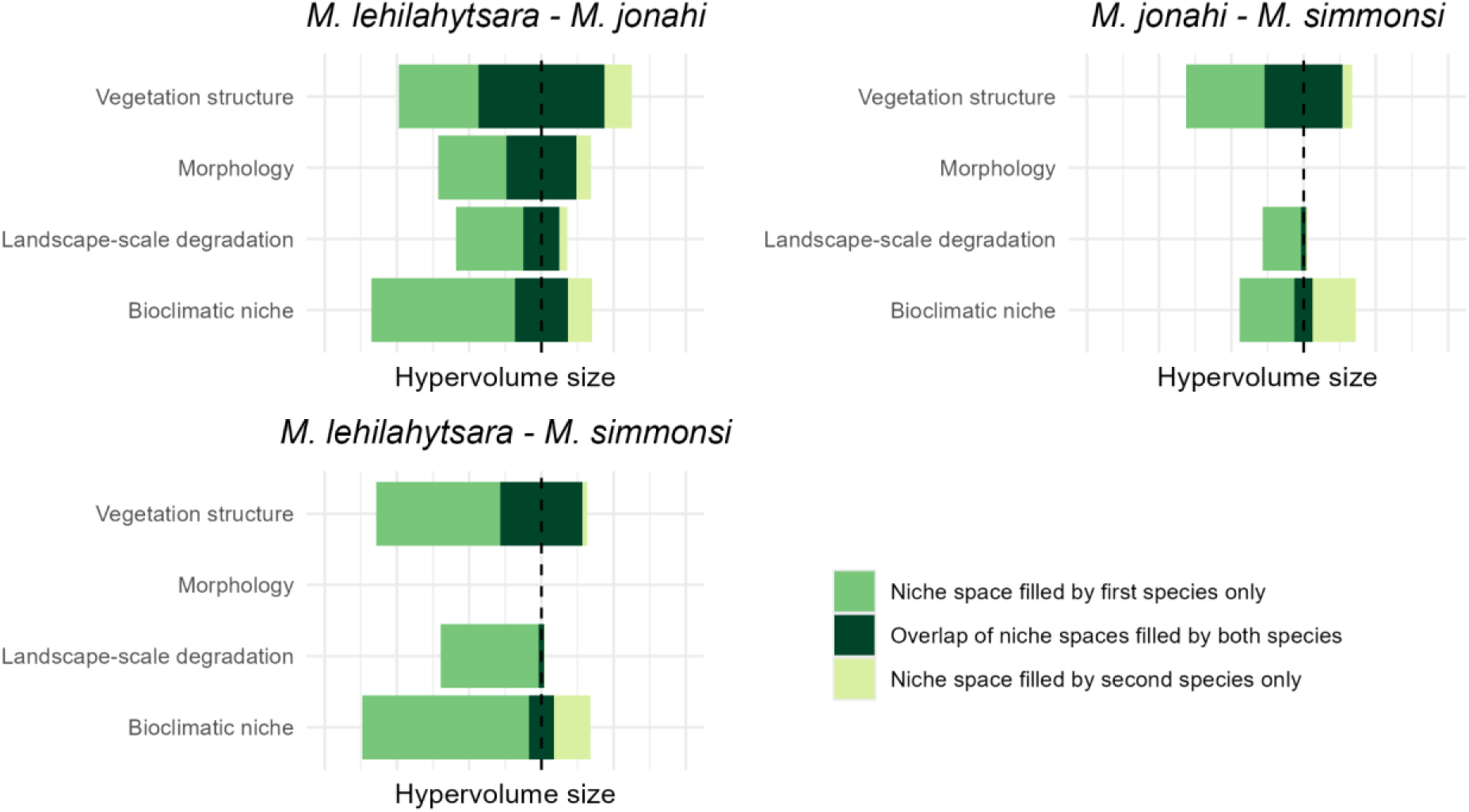
Niche spaces as hypervolume sizes and overlaps between mouse lemur species pairs along multiple trait axes. Sample sizes for niche space estimation were set to the lowest number of records for one species for comparability (vegetation structure n = 62, morphology n = 59 (*M. simmonsi* was excluded due to too few captures), landscape-scale degradation n = 72, bioclimatic niche n = 68). Hypervolume sizes and overlaps are provided in Table S13.

### Annual body mass variability

We observed one clear peak in adult female body mass in *M. jonahi* in November, when all females were pregnant (heaviest female: 115 g). The majority of male *M. jonahi* did not show much annual variability (Figure 5). Noticeably, 17.7% of the 96 individuals captured between March-May, (i.e., before the austral winter) were ≥ 80 g (heaviest female, not pregnant: 118 g; heaviest male, testes regressed: 111 g; outliers in Figure 5). Only these individuals, but not the entire captured population, appeared plump and stored fat throughout their body (Figure S2, S3).

**Figure 5:**
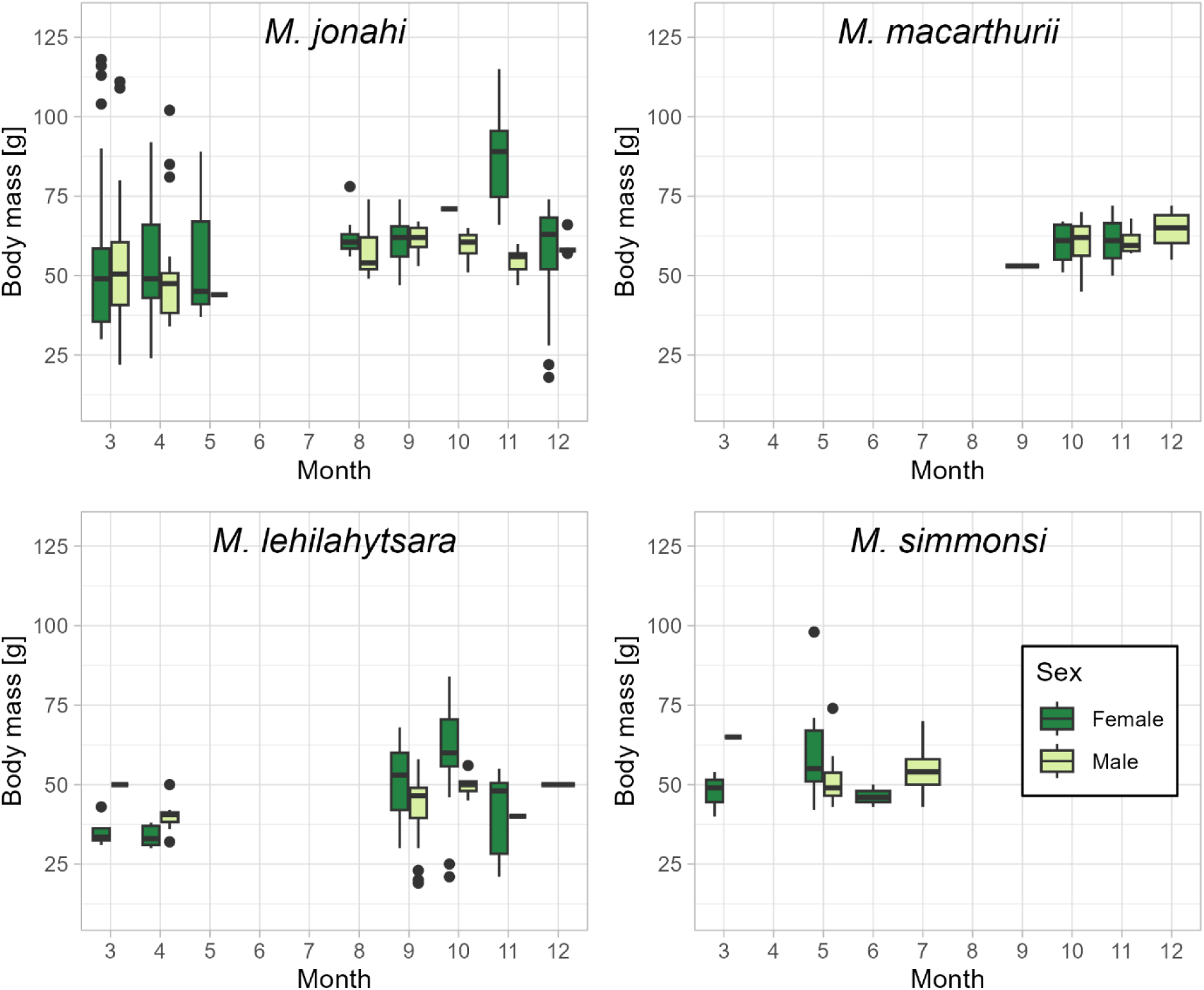
Annual body mass variation in mouse lemurs (*Microcebus* spp.) by month and sex. For photographs refer to Figures S2-S5. Boxplots represent medians with interquartile range (IQR: 25th-75th percentile), whiskers (minimum/maximum values) and outliers (values >1.5*IQR further away from the IQR itself).

For the partly co-distributed *M. lehilahytsara*, the first peak (i.e., weight gain of pregnant females) was also observed (heaviest female: 84 g), although already in October. No second peak with fat individuals was observed, but 38.9% of specimens stored fat in their tails between March-May (n = 18; tail diameter ≥ 0.45 cm; Figure S4, S5).

About 13.0% of the 23 captured *M. simmonsi* in March-May were also very heavy with fat deposits stored throughout the whole body, comparable to *M. jonahi* (heaviest female, not pregnant: 98 g; heaviest male, testes regressed: 74 g, Figure 5). A comparable analysis was not possible for *M. macarthurii*, due to capture data being only available from September-December (n = 20 individuals across 4 months; Figure 5).

## DISCUSSION

### Niche adjustment enables sympatric occurrences

Cryptic and closely related species are generally considered to rarely coexist due to a particularly high overlap in resource use based on their morphological and physiological similarity (Cothran et al. 2013; Violle et al. 2011; Vodā et al., 2015). We therefore expected similar mouse lemur densities in allopatric and sympatric distributions, aligning with the theory of density compensation (MacArthur and Wilson, 1967). However, about 2.4 times more individuals were observed in sympatric areas, with higher encounter rates across all habitat types on average, suggesting a certain degree of niche partitioning.

In general, the four species were found in a variety of habitats, ranging from undisturbed to highly logged primary forests, from agroforestry systems to woody or shrubby fallows. Mouse lemurs were absent from agricultural fields, degraded fernlands and pastures, consistent with the literature (Hending et al., 2018; Knoop et al., 2018; Miller et al., 2018; Webber et al., 2020).

They regularly occurred outside of forests but only within a margin of about 600 m from the forest edge (see supplementary discussion), suggesting that fallow-derived habitats are suitable dispersal matrices but may not allow for stable populations to form, if no forests are nearby (Schüßler et al., 2018). When forests are within reach, lemurs can retreat in response to fallow-to-field cycles (i.e., shifting cultivation), but if isolated, they risk entrapment and local extinction.

*M. jonahi* and *M. simmonsi* were equally often found in forest- and fallow-derived habitats (*M. simmonsi*: Miller et al., 2018), whereas *M. lehilahytsara* was generally more often observed in fallows. This shift was found under sympatric conditions but not in areas of its allopatric occurrence. In allopatry in general, abiotic and biotic constraints like intra-specific competition for critical resources may be the primary factor limiting the population density of mouse lemurs (Brown, 1984; Santini et al., 2023; Stephens et al., 2019). In the case of sympatry, though, a second species appears to be unaffected by the constraints impacting the other species. A limitation of food resources is unlikely in the highly productive humid lowland forests of northeastern Madagascar, as most of the studied mouse lemurs are known to have a flexible and broad omnivorous dietary spectrum, including fruits, leaves and arthropods (Atsalis, 1999; Dammhahn and Kappeler, 2008a; Radespiel et al., 2006; Ramananjato et al., 2020). Limitations of space may be only partly expected, as territoriality in mouse lemurs is not too strong, given overlapping home ranges, scramble competition, the formation of sleeping groups in both sexes and only some degree of intra-sexual agonistic behaviors during nocturnal encounters (Dammhahn and Kappeler, 2008b; Evasoa et al., 2019; Génin, 2008; Jürges et al., 2013; Karanewsky and Wright, 2015; Radespiel, 2000; Schwab and Ganzhorn, 2004). A limited availability of sleeping sites and shelters instead may be more likely. Although we have no data on sleeping site selection for *M. jonahi* and *M. macarthurii*, their association to forest-derived habitats may be indicative for the use of tree holes or wooden shelters as preferred sleeping sites. This was anecdotally noted for *M. simmonsi* on Ile Ste. Marie (Goodman, 1993) and regularly reported for other species like *M. murinus* (Radespiel et al., 2003). In contrast, *M. lehilahytsara* is known to use tree holes but also to build nests out of leaves and branches (Andriambeloson et al., 2021), comparable to *M. rufus*, *M. sambiranensis* and *M. ravelobensis* (Hending et al., 2017; Karanewsky and Wright, 2015; Radespiel et al., 2003; Thorén et al., 2010). This may suggest a higher flexibility to adapt to fallow-derived habitats compared to *M. macarthurii* and *M. jonahi*.

*Microcebus lehilahytsara* exhibited the largest niche breadth across all trait axes. This is consistent with expectations for its bioclimatic niche, given its broader geographic distribution and wide elevational amplitude (up to 1,552 m a.s.l.; Table S14; van Elst et al., 2024; Tiley et al., 2022). Furthermore, its broader niche regarding landscape-scale degradation and habitat structures supports the hypothesis that its extensive distribution is driven by a high ecological plasticity (Kambach et al., 2019; Slatyer et al., 2013). This adaptability is also reflected in its greater morphological variability, aligning with the niche variation hypothesis (Bolnick et al., 2007; Franco-Trecu et al., 2022; Maldonado et al., 2017; van Valen, 1965). It has been suggested before that mouse lemurs may generally exhibit morphological adaptations towards local bioclimatic conditions, for example, following Allen’s rule (Schüßler et al., 2023a).

To summarize, the larger niche spaces of *M. lehilahytsara*, its flexibility in sleeping site and shelter use, together with its shifted occurrence towards fallow-derived habitats in sympatry suggest that this species can adjustment its niche to allow for coexistence with the larger-bodied and closely related *M. jonahi* and *M. macarthurii* (Costa-Pereira et al., 2018). A comparable relationship with one species adjusting its niche to promote coexistence with a second species has also been found in howler monkeys (*Alouatta* spp.; Flores-Escobar et al., 2020). However, why this potential for coexistence is not equally realized in all inter-river systems where *M. lehilahytsara* could co-occur with *M. jonahi,* remains unkown.

### Competition likely maintains allopatry

*M. jonahi* and *M. simmonsi* had smaller niche space sizes compared to *M. lehilahytsara*. We estimated only minor niche overlaps along all trait axes, but also notable niche differentiation that could potentially facilitate coexistence. However, *M. simmonsi* exhibited the smallest niches along all four axes. It was not found in sympatry with any other mouse lemur species, although this would theoretically be possible based on its broad elevational amplitude ranging from sea level to 992 m a.sl., which is comparable to *M. jonahi* (Table S14). Specifically in inter-river system 11 (Figure 1), its absence from mid-elevations, where *M. jonahi* was found instead, is intriguing. However, it appears to be more specialized concerning habitat structures, which may limit its plasticity and likely also its competitive potential (Büchi and Vuilleumier, 2014; Ramiadantsoa et al., 2018). Like *M. jonahi, M. simmonsi* are most often found in forest-derived habitats, which may not enable them to adjust their niches to avoid competition for critical resources like tree holes or wooden shelters. These ecological constraints of *M. simmonsi* may generally prevent its coexistence with other mouse lemurs (Costa-Pereira et al., 2018; Flores-Escobar et al., 2020). We would instead expect a contact zone between *M. jonahi* and *M. simmonsi* somewhere in the lowlands of inter-river system 11. Comparable to a case in mouse opossums (*Marmosa* spp.) in Venezuela (Gutiérrez et al., 2014), allopatry may be maintained by a likely competition for critical resources between *M. jonahi* and *M. simmonsi*. However, this hypothesis requires further investigation.

### Annual body mass variability suggests use of heterothermy

Mouse lemurs represent a basal group of primates with unique features such as the ability to flexibly exhibit heterothermy to withstand unfavorable ambient conditions (Blanco et al., 2016, 2018; Dausmann and Warnecke, 2016). A prerequisite for heterothermy is the ability to store fat for times of inactivity (Bachorec et al., 2023; Sheriff et al., 2013; Vuarin et al., 2015). *M. lehilahytsara*, a species already known to exhibit prolonged torpor (Andriambeloson et al., 2020), apparently stores fat primarily in its tail, leading to an increase in body weight of 40-50% from the onset of the breeding season to the onset of the austral winter (Blanco et al., 2016; Randrianambinina et al., 2003). We also found *M. lehilahytsara* with fattened tails during our study. In addition, we documented *M. jonahi* and *M. simmonsi* individuals with fat deposits throughout their body, not just in the tail. Assuming an average body weight of about 60 g at the start of the reproductive season (October) for both species (Schüßler et al., 2020a, 2023a). Weight gain for *M. jonahi* ranged from 85-97% at the onset of the austral winter (i.e., April, May). Slightly lower but comparable values of weight gain of up to 80% were reported for heterothermic *M. rufus* and *M. murinus* (Atsalis, 1999; Chazarin et al., 2022). Our preliminary data suggests that individuals of both species (*M. jonahi* and *M. simmonsi*) have the prerequisites to exhibit heterothermy.

A larger amount of stored fat has been shown in different mammals to enable a more flexible use of heterothermy, as an adaptation to unpredictable environmental conditions, predator avoidance or reproductive strategies (Allison et al., 2023; Bieber et al., 2013; Zervanos et al., 2013). The use of daily (< 24 h) or prolonged torpor (>24 h) or even hibernation for several weeks was already reported in six of the 19 different mouse lemur species (Blanco et al., 2016, 2018; Dausmann and Warnecke, 2016). Heterothermy is flexibly exhibited in different species, between populations, and even within the same population, depending on an individual’s body condition (Kobbe and Dausmann, 2009; Ortmann et al., 1997; Schmid and Kappeler, 1998). Leaner individuals with less fat deposits may only enter daily torpor, while those with strong gains in body mass may exhibit hibernation (Kobbe and Dausmann, 2009; Ortmann et al., 1997; Schmid and Kappeler, 1998). Interestingly, we found extreme variance of weight gain within populations of *M. jonahi* and *M. simmonsi*, with some individuals not showing any signs of fattening, others having only fattened tails, and again others with dramatic weight gain and fat deposits throughout their body. This high flexibility in fat deposition strategies may be an important prerequisite and selective advantage for the persistence of populations under unpredictable environmental conditions (Blanco et al., 2018; Dausmann, 2014; Dausmann and Warnecke, 2015). Furthermore, species-specific differences in the patterns of heterothermy could provide a basis for coexistence during times of limited resource availability (Adams and Thibault, 2006; Campera et al, 2019; Snodderly et al., 2019). However, the lack of sympatry between *M. jonahi* and *M. simmonsi* in IRS 11 does not support this scenario but rather favors the hypothesis that both species may directly or indirectly compete for sleeping sites that can have essential functions for mouse lemurs regarding predator avoidance, infant rearing but also for thermoregulation during periods of inactivity (Blanco et al., 2016; Lutermann et al., 2010; Karanewsky and Wright, 2015; Thorén et al., 2010).

### Synthesis: Explaining patterns of allopatry and sympatry

Species distributions are shaped by a complex interplay of biotic and abiotic factors, ecological plasticity, evolutionary history, and dispersal abilities (Acevedo et al., 2016; Boulangeat et al., 2012; Gaston 2003; Sexton et al., 2009). While Schüßler et al. (2025) and van Elst et al. (2025) investigated the abiotic constraints and phylogenetic trajectories of mouse lemur phylogeography in northeastern Madagascar, this study integrates their ecological relationships to explain their modern distributions.

Habitat use and niche characteristics could be related to the range size of the investigated mouse lemur species (niche breadth-range size hypothesis; Kambach et al., 2019; Slatyer et al., 2013). A wider distribution across variable bioclimatic conditions, together with flexibility in habitat and likely sleeping site use suggest an elevated ecological plasticity for *M. lehilahytsara*, which allows for niche adjustment in cases of sympatry with the larger *M. jonahi* and *M. macarthurii*. In contrast, *M. jonahi* and *M. simmonsi* are likely to form a parapatric contact zone in which competition for sleeping sites may maintain their allopatry (Anderson, 2016; Gutiérrez et al., 2014). Under these conditions, *M. lehilahytsara* can be classified as an ecological generalist species (Büchi and Vuilleumier, 2014; Ramiadantsoa et al., 2018) compared to the lowland rainforest specialists *M. jonahi*, *M. macarthurii* and *M. simmonsi*.

The elevated ecological plasticity of *M. lehilahytsara* would suggest its presence in all lowland sites where *M. jonahi* or *M. simmonsi* can be found today. However, this was not observed. We hypothesize that the interaction of two factors can explain this pattern: (1) a different capacity to use heterothermy, deduced from differential patterns of fattening between *M. lehilahytsara*, *M. jonahi* and *M. simmonsi* (Andriambeloson et al., 2020; Blanco et al., 2016; this study), and (2) that modern distributions are a relic of historical colonization and habitat suitability (Dausmann and Warnecke, 2015; Nowack and Dausmann, 2016). The capacity to deposit larger amounts of fat may generally provide a higher flexibility to persist under unpredictable environmental conditions (Allison et al., 2023; Bieber et al., 2013; Blanco et al., 2018; Dausmann, 2014; Wright, 1999; Zervanos et al., 2013). The paleoclimatic oscillations of the Pleistocene, during which the diversification of the genus *Microcebus* likely happened (van Elst et al., 2024; Yoder et al., 2016), were characterized by an exceptional unpredictability (Hofreiter and Stewart, 2009; Lupien et al., 2020) During a glacial maximum, increased aridity and lower annual mean temperatures in the tropics have led to a contraction of forests to areas still receiving sufficient water supply (Gasse and Van Campo, 2001; Teixeira et al., 2021; Wilmé et al., 2006). These were likely located in mountain refugia, where orographic precipitation was captured, or in lowland riparian wetlands at topographically advantageous locations (Duminil et al., 2015; Everson et al., 2020; Schüßler et al., 2025; Mercier and Wilmé et al., 2013). If we expect the latter to be less predictable, due to the dependence on orographic precipitation and the downstream flow of surface water towards the lowlands, it can be argued, that increased heterothermic flexibility allowed for persistence in the more unpredictable lowlands, too (Nowack and Dausmann, 2015). Following this argument, the high capacity of fat storage observed in *M. jonahi* may have allowed this species to colonize and survive in all inter-river systems (IRS 6-14) while *M. lehilahytsara* may have retreated from or gone extinct in lowland areas it inhabited before the most arid and therefore challenging times of the last glacial maximum. This could also explain why the small-bodied *M. lehilahytsara* was not yet found in sympatry with the other large-bodied lowland species, *M. simmonsi* and *M. gerpi*, although their ranges would be in reach and the ecological plasticity to adjust its niche has been documented. Further detailed ecological studies are still required to test the hypotheses generated in this study.

## IMPLICATIONS FOR CONSERVATION

Mouse lemurs, like many other species (e.g., birds, amphibians, reptiles, endemic herbaceous plants or endemic arthropods; Fulgence et al., 2021; Martin et al., 2020a; Rakotomalala et al., 2021; Raveloaritiana et al., 2021; Wurz et al., 2022) are bound to forest habitats. Although presences are often reported in fallow-derived habitats (Hending et al., 2018; Knoop et al., 2018; Webber et al., 2020), the observed strong decline of sightings with increasing distance from the forest edge suggests that non-forest habitats may mainly serve as dispersal or foraging matrices and may act as population sinks if situated too far from forests. Given widespread deforestation in eastern Madagascar, fallows should not be considered suitable habitats unless nearby forest fragments persist. This applies to IUCN-listed Endangered species like *M. gerpi*, *M. jollyae, M. jonahi, M. macarthurii and M. simmonsi*.

A major implication of this study and that of van Elst et al. (2025) is that distributions can be unexpectedly disjunct (i.e., lowland absences of *M. lehilahytsara*, absence of *M. simmonsi* from IRS 12-14), which were previously thought to be continuous based on extrapolation from fewer presence records and elevational ranges. *Microcebus* represents the best sampled vertebrate genus from the northern half of the Malagasy east coast, allowing for a comprehensive investigation of the patterns of evolution, diversity and distributions (van Elst et al., 2023, 2024, 2025). Whether comparable mechanisms and interactions also modulate distributions of other taxa remains elusive today. Unfortunately, the time for uncovering evolutionary patterns and the true biodiversity of eastern Madagascar is running out: lowland rainforests were cleared in south- and central-eastern Madagascar already 50+ years ago (Vieilledent et al., 2018), while northeastern Madagascar has lost more than 20% of its forest cover since 1990 (Schüßler et al., 2020b). Together with the projected ongoing rate of deforestation (Morelli et al., 2019; Schüßler et al., 2020b), our findings call for an increase in conservation action and research on other taxa, before it is too late.

## ACKNOWLEDGEMENTS

We sincerely thank the Malagasy authorities for granting us permission to conduct this research. These are the Ministry of Environment and Sustainable Development (formerly the Ministry of Environment, Water and Forests), the Direction des Aires Protégées, des Réssources Naturelles Renouvelables et des Ecosystemes, its regional authorities (DREDD), the heads of the Cantonnement and the local village leaders who allowed us to visit their forests. This research was conducted under the permits 197/17/MEEF/SG/DGF/DSAP/SCB.Re, 169/19/MEDD/SG/DGEF/DGRNE, 349/21/MEDD/SG/DGGE/DAPRNE/SCBE.Re, 030/22/MEDD/SG/DGGE/DAPRNE/SCBE.Re. We are grateful to Dr. Brigitte M. Raharivololona, Prof. Jonah Ratsimbazafy and GERP (Groupe d’Etude et de Recherche sur les Primates) for their support in project coordination and infrastructure. We are further thankful for the help during data collection of Yves Rostant Andriamalala, Maria Jäger, Deborah Rabesamihanta, Ange Nandrianina, Fils Donna Rakotoarizaka, Editho Bellarmain Beandakana, and our local guides, cooks, and porters. Funding was kindly provided by German Science Foundation (DFG, RA 502/23-1) to U. R., the Society for Tropical Ecology, Houston Zoo, the Bauer Foundation (T237/22985/2012/kg) and the Zemplin Foundation (T0214/32083/2018/sm) to D. S.

## AUTHOR CONTRIBUTIONS

U.R., D.S and J.M.C. conceived the study; D.S., T.v.E., N.R.R., T.R. S.M.R., R.D.R, and E.R. collected the data; D.S. analyzed the data; D.S. wrote the first draft. All authors contributed to editing drafts and gave final approval to submit the manuscript.

