## Supplementary Information for "Ecological plasticity explains the distribution of sympatric and allopatric mouse lemurs (*Microcebus* spp.) in northeastern Madagascar"

#### Supplementary methods

##### Mouse lemur occurrences in fallow-derived habitats

The number of transects per habitat type and under sympatric and allopatric occurrences is summarized in Table S1.

Additionally, for all mouse lemur sightings outside of forest-derived habitats, we estimated the Euclidean aerial distances between the sighting locations to the nearest forest in QGIS as derived from the land cover maps of Schüßler et al. (2020b). Our transects ranged in distance from the forest edge from 0-7,710 m.

**Table S1:** Number of transects and survey effort across the different habitat types and for allopatric and sympatric sites of the study region.

| | | **Allopatric sites** | | **Sympatric sites** | | **Total** | |
|---|---|---|---|---|---|---|---|
| **Habitat type** | | Number of transects | Survey effort [km] | Number of transects | Survey effort [km] | Number of transects | Survey effort [km] |
| Undisturbed primary forest | PF | 11 | 32.539 | 4 | 7.331 | 15 | 39.870 |
| Selectively logged primary forest | SLPF | 21 | 72.809 | 9 | 25.260 | 30 | 98.070 |
| Highly logged primary forest | HLPF | 22 | 51.954 | 12 | 30.068 | 34 | 82.023 |
| Agroforestry | AF | 3 | 10.926 | 4 | 4.977 | 7 | 15.903 |
| Fallow land | Savoka | 14 | 37.960 | 4 | 10.225 | 18 | 48.185 |
| Fernland | Fernland | 5 | 26.387 | 0 | 0.000 | 5 | 26.387 |
| Sum | | 76 | 232.576 | 33 | 77.861 | 109 | 310.437 |

**Table S2:** Variance explained by the principal components used for the estimation of hypervolumes across four different environmental and phenotypic axes.

| | PC1 | PC2 | PC3 | PC4 | PC5 | PC6 | PC7 | PC8 | PC9 |
|---|---|---|---|---|---|---|---|---|---|
| **Vegetation structure** | | | | | | | | | |
| Standard deviation | 1.7153 | 1.2501 | 1.0251 | 0.89013 | 0.72960 | 0.69446 | 0.64131 | 0.47547 | |
| Proportion of variance | 0.3678 | 0.1953 | 0.1313 | 0.09904 | 0.06654 | 0.06028 | 0.05141 | 0.02826 | |
| Cumulative variance | 0.3678 | 0.5631 | 0.6945 | 0.79351 | 0.86005 | 0.92033 | 0.97174 | 1.00000 | |
| **Landscape-scale degradation** | | | | | | | | | |
| Standard deviation | 1.2766 | 1.0993 | 0.9083 | 0.58810 | | | | | |
| Proportion of variance | 0.4065 | 0.3014 | 0.2058 | 0.08627 | | | | | |
| Cumulative variance | 0.4065 | 0.7079 | 0.9137 | 1.00000 | | | | | |
| **Bioclimatic niche** | | | | | | | | | |
| Standard deviation | 2.2502 | 1.4155 | 1.1174 | 0.8234 | 0.55622 | 0.19528 | 0.16315 | 0.07868 | 0.06749 |
| Proportion of variance | 0.5399 | 0.2136 | 0.1331 | 0.0723 | 0.03299 | 0.00407 | 0.00284 | 0.00066 | 0.00049 |
| Cumulative variance | 0.5399 | 0.7535 | 0.8867 | 0.9590 | 0.99195 | 0.99602 | 0.99885 | 0.99951 | 1.00000 |
| **Morphology** | | | | | | | | | |
| Standard deviation | 1.6184 | 1.0918 | 0.9281 | 0.72217 | 0.67590 | 0.59058 | | | |
| Proportion of variance | 0.4366 | 0.1987 | 0.1436 | 0.08692 | 0.07614 | 0.05813 | | | |
| Cumulative variance | 0.4366 | 0.6352 | 0.7788 | 0.86573 | 0.94187 | 1.00000 | | | |

**Table S3:** Variable contributions to the first three principal components (PC) used for estimating the hypervolumes of vegetation structure.

| Variable | PC1 | PC2 | PC3 |
|---|---|---|---|
| Coverage < 0.5 m | 0.4083 | 0.2888 | -0.0458 |
| Coverage 0.5.2 m | 0.4328 | 0.3972 | -0.0649 |
| Coverage 2-5 m | 0.2173 | 0.5648 | -0.0586 |
| Coverage 5-10 m | -0.3261 | 0.4476 | -0.3443 |
| Coverage > 10 m | -0.4151 | 0.2791 | -0.1830 |
| Coverage of *Harungana madagascariensis* | 0.2697 | -0.1891 | -0.6996 |
| Coverage of *Aframomum angustifolium* | 0.3465 | 0.0702 | 0.4988 |
| Coverage of *Clidemia hirta* | 0.3568 | -0.3476 | -0.3162 |

**Table S4:** Variable contributions to the first three principal components (PC) used for estimating the hypervolumes of morphology.

| Variable | PC1 | PC2 | PC3 |
|---|---|---|---|
| Ear length | 0.0507 | 0.7901 | -0.4246 |
| Head length | -0.4288 | 0.0955 | -0.5354 |
| Head width | -0.4199 | 0.2265 | 0.5533 |
| Interorbital distance | -0.4700 | 0.2925 | 0.2590 |
| Body length | -0.5274 | -0.1146 | 0.0048 |
| Tail length | -0.3717 | -0.4655 | -0.3998 |

**Table S5:** Variable contributions to the first three principal components (PC) used for estimating the hypervolumes of landscape-scale degradation.

| Variable | PC1 | PC2 | PC3 |
|---|---|---|---|
| Population density | -0.7116 | -0.0458 | 0.0763 |
| Distance to nearest village | -0.3598 | 0.6095 | 0.5880 |
| Forest density | 0.6034 | 0.3097 | 0.4402 |
| Deforestation density | 0.0002 | -0.7284 | 0.6743 |

**Table S6:** Variable contributions to the first three principal components (PC) used for estimating the hypervolumes of bioclimatic niche.

| Variable | PC1 | PC2 | PC3 |
|---|---|---|---|
| Elevation | -0.3941 | 0.2850 | 0.0848 |
| Isothermality | -0.3358 | -0.1271 | -0.5078 |
| Temperature seasonality | -0.2937 | 0.2243 | -0.5331 |
| Maximum temperature of warmest month | 0.2663 | -0.3794 | -0.4274 |
| Minimum temperature of coldest month | 0.4066 | -0.2692 | 0.1895 |
| Annual precipitation | 0.3622 | 0.4244 | -0.0911 |
| Precipitation seasonality | -0.3512 | 0.0436 | 0.3474 |
| Precipitation of wettest quarter | 0.1512 | 0.5896 | 0.0762 |
| Precipitation of driest quarter | 0.3634 | 0.3259 | -0.3120 |

#### Supplementary results

The population density of *Microcebus jonahi* in allopatry over all habitat types was estimated with 78.9 ind./km² (standard error = 12.0, 95% confidence interval: 58.2-106.9 ind./km²; Table S7). Density estimates were not possible for the other species in allopatry due to low sample sizes.

For the mean encounter rates provided in Figure 2, a sampling bias due to seasonality or better visibility could be ruled out, because of an equal sampling of habitats across the study months (not shown) and equally distributed perpendicular distances in all habitats (not shown).

**Table S7:** Population density estimates in individuals per km² for *Microcebus* occurences in sympatry and in allopatry in general and *Microcebus jonahi* in allopatry. Abbreviations: n: sample size; k: number of sampled transects; ind./km²: individuals per square kilometer.

| | **Covered Area [km²]** | **Effort [km]** | **n** | **k** | **Density estimate [ind./km²]** | **Standard error [ind./km²]** | **Coefficient of variation** | **95% confidence interval [ind./km²]** |
|---|---|---|---|---|---|---|---|---|
| Allopatry | 12.035 | 171.936 | 220 | 54 | 72.47 | 8.79 | 0.12 | 56.91 – 92.30 |
| Sympatry | 5.436 | 77.661 | 240 | 32 | 175.04 | 16.29 | 0.09 | 145.06 – 211.21 |
| *M. jonahi* in allopatry | 5.308 | 132.962 | 174 | 39 | 78.91 | 12.01 | 0.15 | 58.23 – 106.93 |

**Table S8:** Tukey post-hoc test after ANOVA for differences in mean encounter rates between habitat types. Abbreviations: 95% CI: 95% confidence interval; P: adjusted P-value after multiple comparisons; PF: undisturbed primary forest; SLPF: selectively logged primary forest; HLPF: highly logged primary forest; AF: agroforestry; savoka: fallow land.

| **Test case** | **Mean difference between groups** | **95% confidence interval** | **P** |
|---|---|---|---|
| PF - AF | -3.02 | -5.86 – -0.18 | 0.03 |
| PF - HLPF | -1.93 | -3.85 – 0.00 | 0.05 |
| SLPF - AF | -2.60 | -5.21 – 0.01 | 0.05 |
| Savoka - AF | -2.72 | -5.48 – 0.05 | 0.06 |
| SLPF - HLPF | -1.51 | -3.06 – 0.05 | 0.06 |
| Savoka - HLPF | -1.62 | -3.43 – 0.19 | 0.10 |
| HLPF - AF | -1.09 | -3.67 – 1.48 | 0.76 |
| SLPF - PF | 0.42 | -1.54 – 2.38 | 0.98 |
| Savoka - PF | 0.31 | -1.86 – 2.48 | 0.99 |
| SLPF - Savoka | 0.11 | -1.74 – 1.97 | 1.00 |

**Table S9:** T-tests on the differences in mean encounter rates between areas of allopatric and sympatric occurrence of mouse lemurs. Abbreviations: t: t-value for test statistic; df: degrees of freedom; P: P-value; PF: undisturbed primary forest; SLPF: selectively logged primary forest; HLPF: highly logged primary forest; AF: agroforestry; savoka: fallow land.

| Habitat type | Mean encounter rate in allopatry | Mean encounter rate in sympatry | t | df | P |
|---|---|---|---|---|---|
| PF | 0.79 ± 0.59 | 1.95 ± 1.75 | -1.300 | 3.251 | 0.280 |
| HLPF | 2.57 ± 3.47 | 3.85 ± 2.00 | -1.360 | 31.830 | 0.183 |
| SLPF | 0.81 ± 0.91 | 3.16 ± 1.86 | -3.608 | 9.700 | 0.005 |
| AF | 1.39 ± 0.65 | 6.16 ± 2.13 | NA | NA | NA |
| Savoka | 1.12 ± 1.30 | 2.40 ± 1.64 | -1.441 | 4.142 | 0.221 |

**Table S10:** Contingency table and results of Poisson test for *Microcebus* spp. occurrences in forest-derived (primary forest (PF), selectively logged primary forest (SLPF), highly logged primary forest (HLPF) versus fallow-derived (agroforestry (AF), fallow with secondary trees (savoka)) habitats. No testing for *M. macarthurii* due to limited sample size.

| # | Species | Forest-derived habitats | Fallow-derived habitats | Rate ratio | P-value |
|---|---|---|---|---|---|
| 1 | *M. jonahi* | 168 | 156 | 1.077 | 0.541 |
| 2 | *M. lehilahytsara* | 18 | 44 | 0.409 | 0.001 |
| 3 | *M. simmonsi* | 42 | 31 | 1.355 | 0.242 |
| 4 | *M. macarthurii* | 1 | 4 | NA | NA |

**Table S11:** Fisher's exact test for *Microcebus* occurrences in forest-derived (primary forest (PF), selectively logged primary forest (SLPF), highly logged primary forest (HLPF) versus fallow-derived (agroforestry (AF), fallow with secondary trees (savoka)) habitats. Occurrence proportions are found in Table S8. No testing for *M. macarthurii* due to limited sample size. Bonferroni-corrected significance level of α = 0.0167 is applied.

| Species 1 | Species 2 | Odds ratio (95% confidence interval) | P-value |
|---|---|---|---|
| *M. lehilahytsara* | *M. jonahi* | 0.381 (0.198-0.706) | 0.001 |
| *M. lehilahytsara* | *M. simmonsi* | 0.305 (0.138-0.655) | 0.001 |
| *M. jonahi* | *M. simmonsi* | 0.800 (0.459-0.137) | 0.437 |

**Table S12:** Contingency table and results of Fisher's exact test for *Microcebus* occurrences in sympatry compared to occurrences in allopatry in forest-derived (primary forest (PF), selectively logged primary forest (SLPF), highly logged primary forest (HLPF) and fallow-derived (agroforestry (AF), fallow with secondary trees (savoka)) habitats. Significant values are given in bold.

| | ***M. jonahi*** | | ***M. lehilahytsara*** | |
|---|---|---|---|---|
| | **Forest-derived habitats** | **Fallow-derived habitats** | **Forest-derived habitats** | **Fallow-derived habitats** |
| Allopatry | 163 | 152 | 12 | 12 |
| Sympatry | 19 | 23 | 6 | 32 |
| Odds ratio | 0.771 | | **0.193** | |
| P | 0.512 | | **0.009** | |

Although fallow-derived habitats appeared to be generally suitable for all mouse lemur species investigated, only occasional sightings occurred with increasing distance from the forest edge (Figure S1). There were continuous sightings of mouse lemurs up to 600 m from the forest edge, but further away only two additional individuals were sighted (at 1,046 m and 1,164 m). The surveyed distances ranged from the forest edge to 7,710 m away from it (Figure S1).

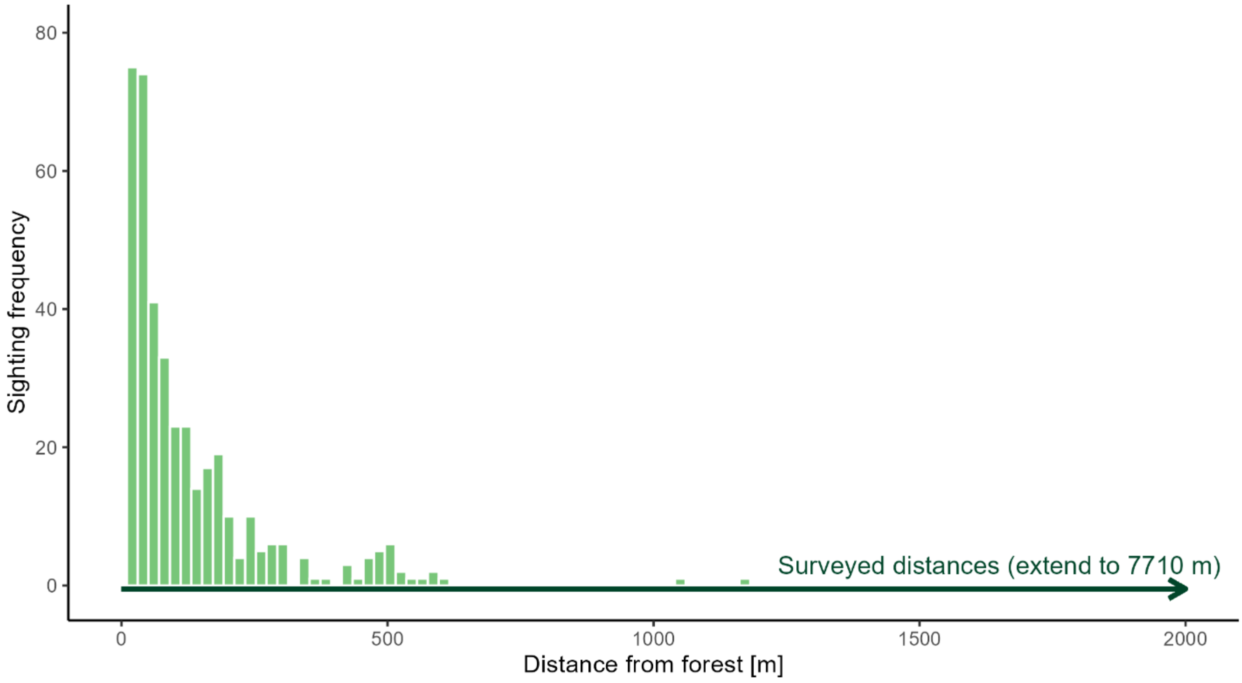


**Figure S1:** Sighting frequencies for mouse lemurs depending on their distance from the forest.

**Table S13:** Hypervolume sizes of *Microcebus* spp. in four different niche dimensions.

| Dimension | Species | n | Hypervolume size | Overlap to *M. lehilahytsara* | Overlap to *M. jonahi* | Overlap to *M. simmonsi* |
|---|---|---|---|---|---|---|
| Vegetation | *M. lehilahytsara* | 62 | 142.65 | | 61.3% | 39.8% |
| | *M. jonahi* | 62 | 106.40 | 82.2% | | 49.6% |
| | *M. simmonsi* | 62 | 60.05 | 94.5% | 88.6% | |
| Landscape | *M. lehilahytsara* | 72 | 71.79 | | 34.8% | 5.3% |
| | *M. jonahi* | 72 | 29.88 | 82.2% | | 11.6% |
| | *M. simmonsi* | 72 | 4.30 | 88.6% | 80.4% | |
| Bioclimatic | *M. lehilahytsara* | 68 | 132.53 | | 27.0% | 12.8% |
| | *M. jonahi* | 68 | 50.74 | 68.5% | | 24.6% |
| | *M. simmonsi* | 68 | 42.34 | 40.1% | 29.5% | |
| Morphology | *M. lehilahytsara* | 59 | 95.82 | | 50.9% | NA |
| | *M. jonahi* | 59 | 58.90 | 82.8% | | NA |
| | *M. simmonsi* | NA | NA | NA | NA | |

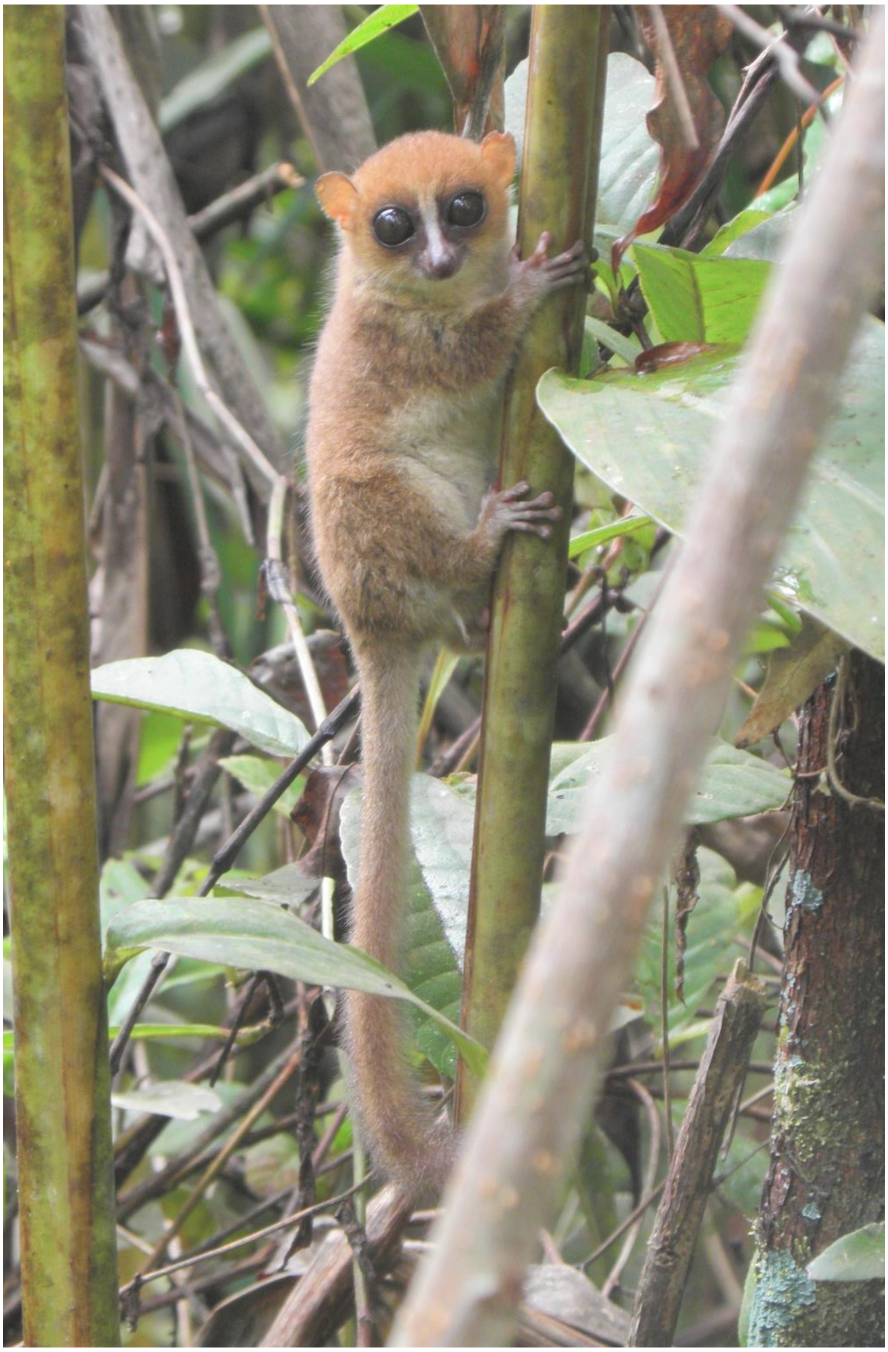

**Figure S2:** Slim *Microcebus jonahi* without fat deposits (captured on 25.03.2022 at an elevation of 639 m a.s.l.) for comparison with Figure S3.

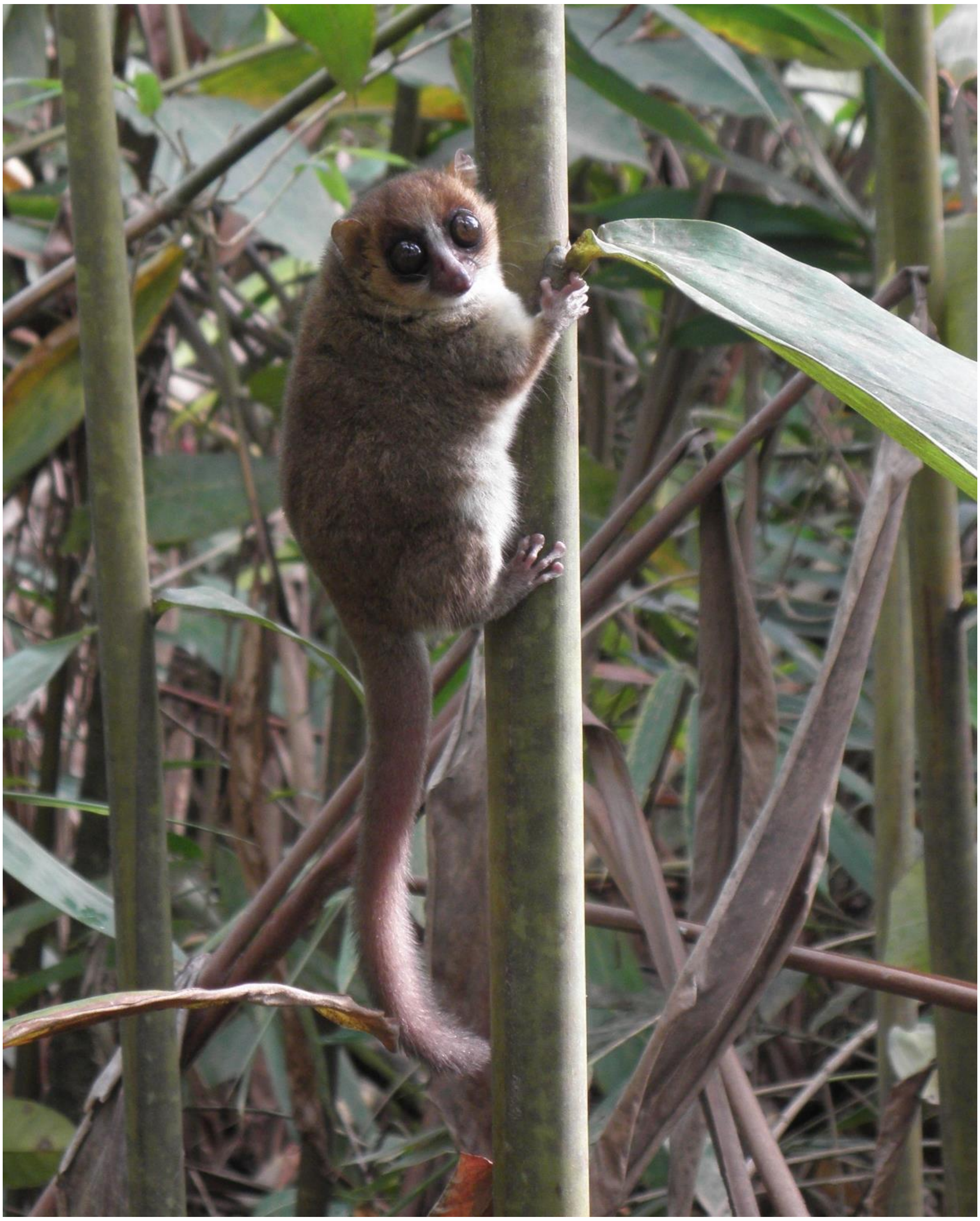

**Figure S3:** Fat *Microcebus jonahi* with fat deposits throughout its entire body (captured on 25.03.2022 at an elevation of 644 m a.s.l.).

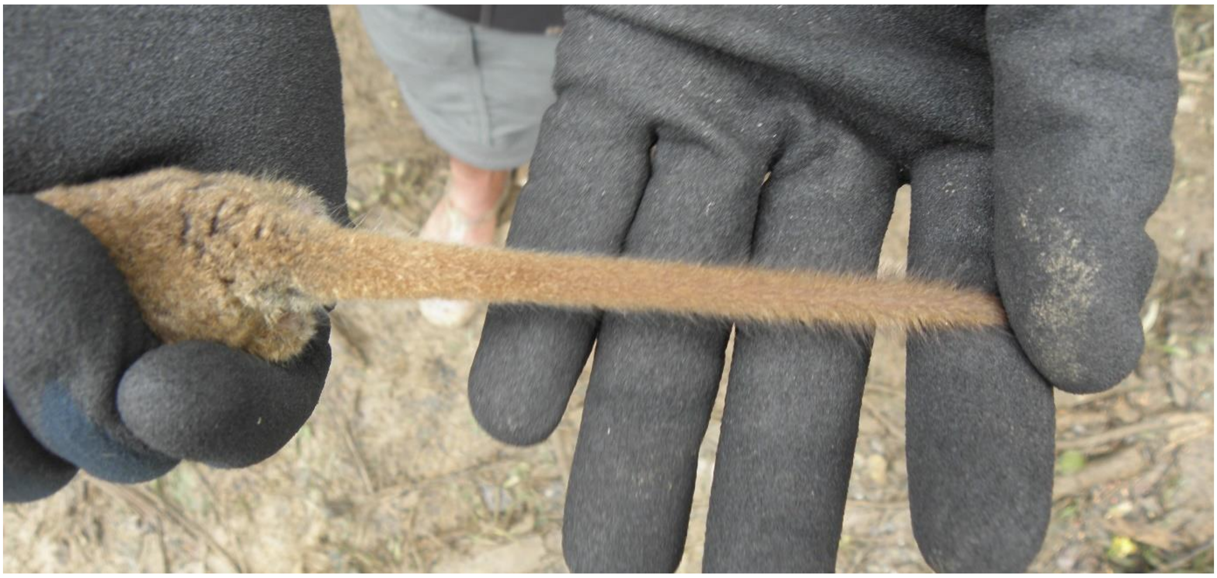

**Figure S4:** Slim tail of *Microcebus lehilahytsara* without fat deposits (captured on 25.03.2022 at an elevation of 637 m a.s.l.) for comparison with Figure S5.

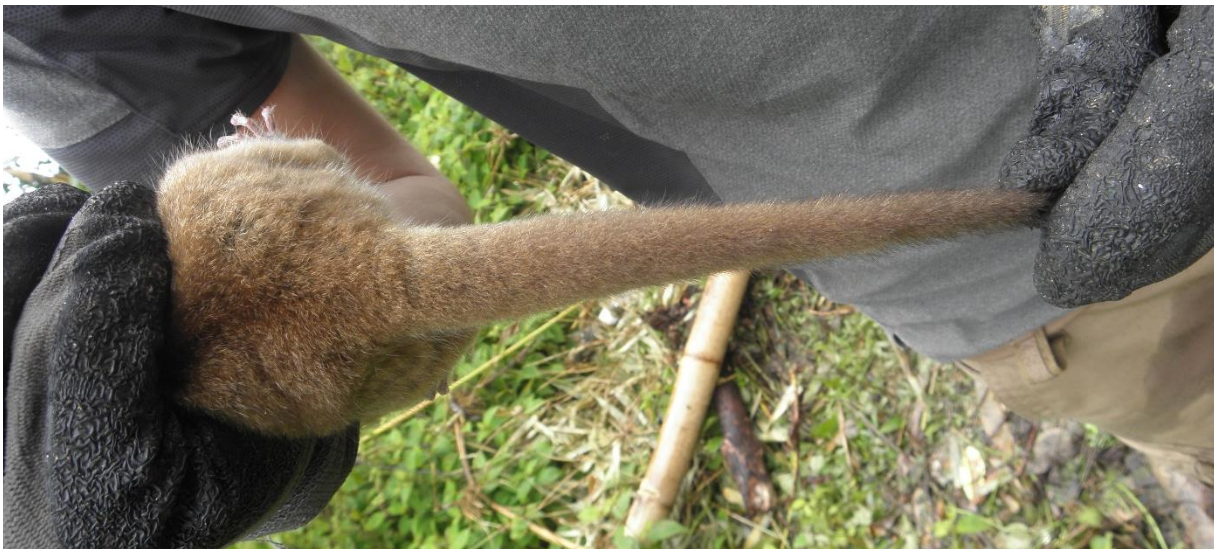

**Figure S5:** Fattened tail of *Microcebus lehilahytsara* (captured on 23.03.2022 at an elevation of 651 m a.s.l.)

### Supplementary discussion

#### Mouse lemur occurrences across different habitat types

The highest abundance of mouse lemurs was found in highly logged primary forests and agroforestry systems. These two habitat types are characterized by the presence of trees, a patchy canopy coverage, a diverse vertical structure and a high abundance of light-dependent plant species. It is not unlikely that these habitats host higher abundances of arthropods (e.g., Malabika, 2011; Perry et al., 2016; Wurz et al., 2022), which could be highly profitable for these small-bodied omnivorous primates (e.g., Lehman et al., 2006; Steffens & Lehman, 2016). Furthermore, a fruit regularly consumed by mouse lemurs is produced by the light-dependent, widespread and invasive *Clidemia hirta* (synonym: *Miconia crenata*), which is present in disturbed habitats and usually found in high abundances. Apart from food resources in HLPF and AF, the presence of trees would also allow for shelter and sleeping sites, which are critical resources for many mouse lemur species (e.g., Blanco et al., 2016; Karanewsky & Wright, 2015; Radespiel et al., 2003).

#### Mouse lemur occurrence in relation to distance from forest edge

Mouse lemur presence in seemingly suitable fallow-derived habitats declined with an increasing distance from the forest edge. Beyond a threshold of about 600 m away from the next forest border, only two mouse lemurs were sporadically sighted, indicating that these habitats are maybe just temporarily used for dispersal or foraging (Schüßler et al., 2018) or, if too far from the forest edge, act as population sinks. If forests are still in reach, this would allow mouse lemurs to retreat or flexibly respond in view of the cyclic transformation of fallows back into agricultural fields, which means that the entire vegetation would be cut down and burnt again. If no suitable habitats were in reach for mouse lemurs inhabiting these areas when the cutting happens, they inevitably would be trapped and go locally extinct. Interestingly, it is often reported that mouse lemurs are captured in fallows close to forests when being prepared for consecutive burning (own data, unpublished). In contrast to that, agroforestry plantations with trees like clove (*Syzygium aromaticum*) or cocoa (*Theobroma cacao*), may buffer against this negative effect by providing shelter and nesting sites for mouse lemurs (Hending et al., 2018; Webber et al., 2020). As wide areas of eastern Madagascar have been deforested and transformed into fallows, agroforestry systems or agricultural fields (Vieilledent et al., 2018; Zähringer et al., 2015), we argue, that these areas should not be considered suitable habitats, if no forest fragment as population source remains in the area. This consideration has major implications for the conservation and estimation of remaining habitat for already Endangered or Critically Endangered species like *M. gerpi* or *M. jollyae*.

#### Geographic distributions of mouse lemurs

Table S14 summarizes information on elevational ranges of the four investigated mouse lemur species and their occurrence per inter-river system (Figure 1)

**Table S14:** Elevational ranges and presence in inter-river-systems (IRS) of the four mouse lemur species (*Microcebus* spp.) of northeastern Madagascar. Numbers of inter-river systems correspond to those given in Figure 1.

| Species | Elevational range [m a.s.l.] | Presence in IRS | Reference |
|---|---|---|---|
| *M. lehilahytsara* | 25 - 1,552 | 2 - 6, 9 - 11 | Andriambeloson et al., 2021; Louis et al., 2006; Poelstra et al., 2021; Radespiel et al., 2008; Rasolofoson et al., 2007; Schüßler et al., 2020a; Weisrock et al., 2010; this study |
| *M. jonahi* | 9 - 953 | 6 - 14 | Louis & Lei, 2016; Poelstra et al., 2021; Schüßler et al., 2020a; this study |
| *M. macarthurii* | 24 - 425 | 5 | Louis & Lei, 2016; Poelstra et al., 2021; Radespiel et al., 2008; Schüßler et al., 2020a; this study |
| *M. simmonsi* | 4 - 992 | 11, 15 - 17 | Louis et al., 2006; Poelstra et al., 2021; Rakotondravony & Rabenandrasana, 2011; Raxworthy, 1986; Schüßler et al., 2020a; this study |
